# Prefrontal orchestration: a cortical network for rodent motor inhibition

**DOI:** 10.1101/2024.10.16.618207

**Authors:** Zoe Jäckel, Niels Schwaderlapp, Ahmed Adzemovic, Florian Steenbergen, Stefanie Hardung, Katharina Fuchs, Christian Leibold, Maxim Zaitsev, Ilka Diester

## Abstract

Goal-directed action control and behavioral flexibility are prerequisites for effective, adaptive behavior. Both abilities rely on functional motor inhibition, which is linked to the prefrontal cortex (PFC), where distinct subsections collaborate in functional networks. How these PFC subsections interact and which roles they play during motor inhibition remains incompletely understood. In this study, we employed an action-preparation task in rats, combined with bidirectional optogenetic interventions, opto-fMRI, single unit electrophysiology and local field potential synchrony measurements across PFC subsections. Our findings support a clear and simple model of action inhibition within the prefrontal network. This model suggests prelimbic cortex (PL) as an input-dependent switch between motor inhibition and execution, modulated by an infralimbic cortex (IL)-dominated network. This distribution of tasks allows the PL to mediate goal-directed action while the IL ensures behavioral flexibility.

Graphical Abstract:
Behavioral measurements were conducted alongside optogenetic modulation of PL, IL or VO. Inhibitory modulation led to varying effects on performance, while excitatory ChR2 stimulation of PL, IL or VO led to analogous effects on proactive motor inhibition. To identify shared nodes recruited by ChR2 stimulation of distinct PFC subareas, we performed whole-brain mapping with opto-fMRI. This revealed an overlapping activation volume spanning PFC, BF, Fr, Cg2, and M2. Notably, this common activation volume closely outlined the entirety of the IL-recruited regions; IL excitation also produced robust behavioral effects. Multisite recordings revealed task performance-dependent PL-IL delta synchrony. PCA of single-unit activity during behavior revealed varied neural patterns among PFC subsections, highlighting PL to have the most the homogenous input-driven activity. The findings can be interpreted as PL acting as an input-dependent switch between motor inhibition and execution, modulated by IL to maintain behavioral flexibility.

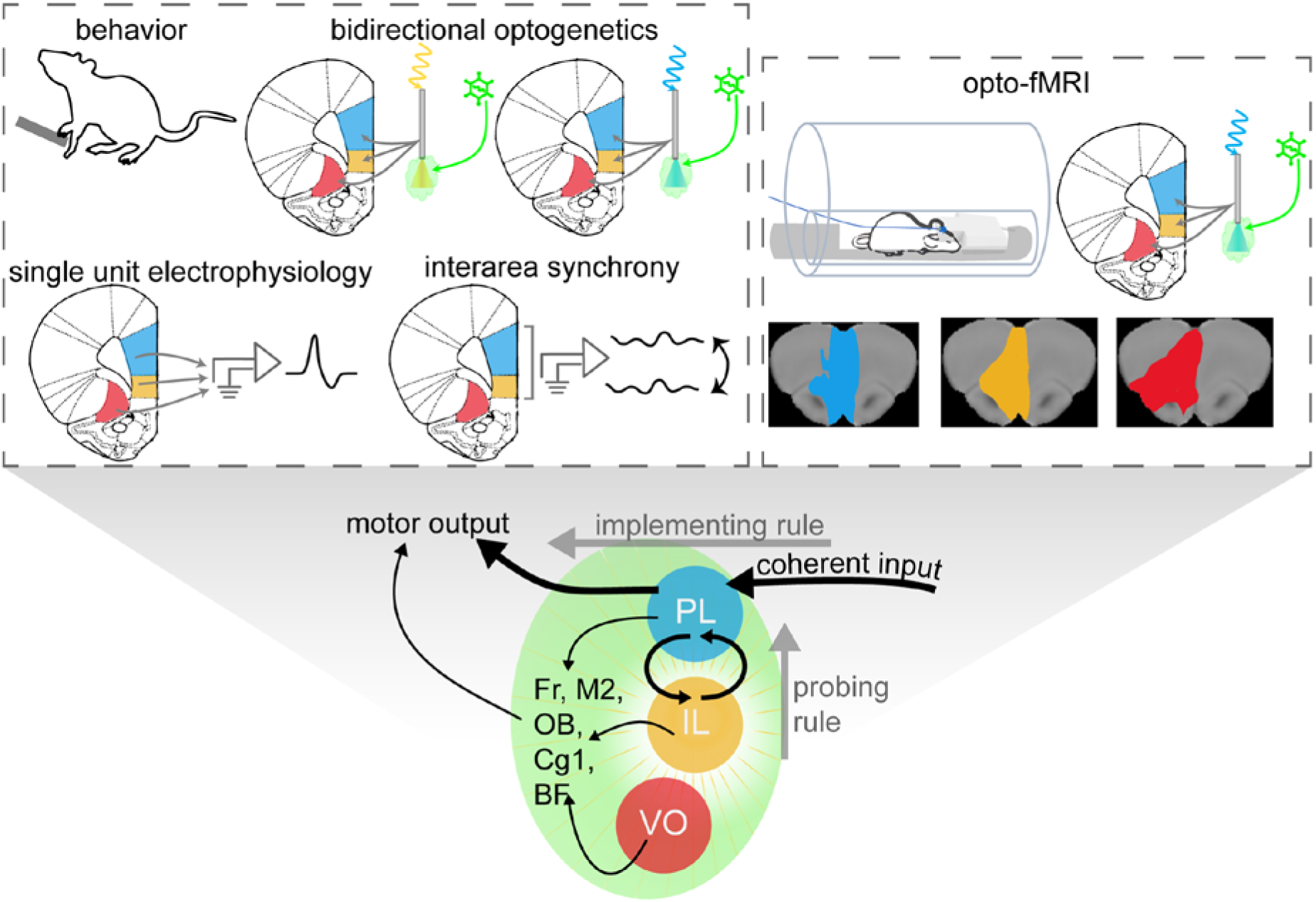

## Introduction

Goal-directed action control^1,2^ and adaptive behavior^3^ rely on the prefrontal cortex (PFC), where distinct subsections collaborate in functional networks^4^. The extent to which this network exhibits a hierarchical system, characterized by early convergence within the PFC, remains elusive^5^. Based on subsection-specific optogenetic inhibition, we have previously described distinct roles of rat PFC subsections in action inhibition with prelimbic cortex (PL) holding back premature responses, infralimbic cortex (IL) promoting these premature responses, and ventro-orbital cortex (VO) being involved in reactive responses^4^. Given that the PL is the most relevant subsection relating to mPFC motor control^1,6^, its role in motor inhibition is intuitive. We thus set out here, to test the hypothesis that PL excitation would lead to an opposing effect to PL inhibition. Indeed, we could confirm this hypothesis (Fig. 1). However, IL and VO excitation led to surprisingly similar effects to PL excitation, which prompted the here described series of experiments.

**Figure 1:**
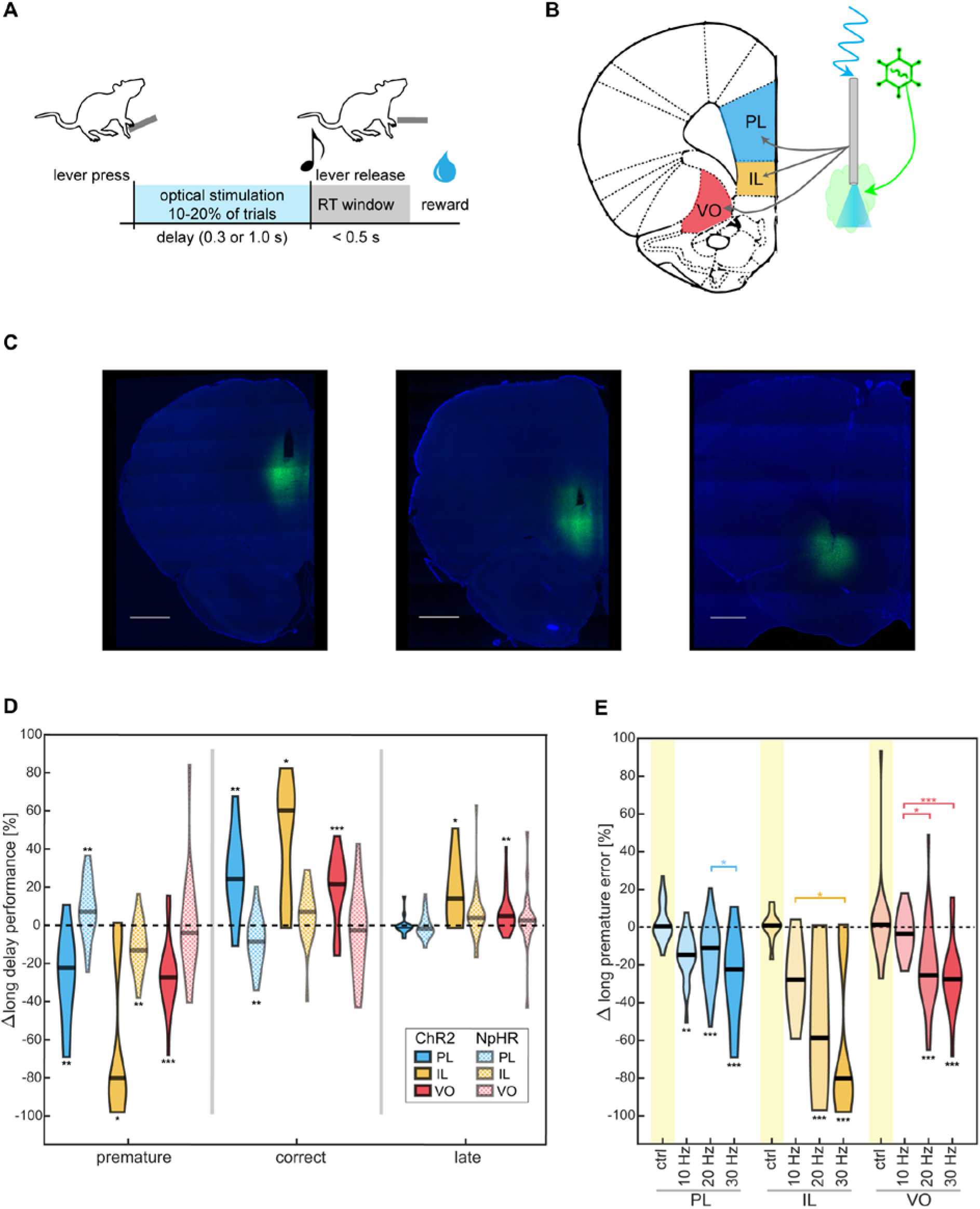
Optogenetic perturbation during motor inhibition. **(A)** Response-preparation task; adapted from Hardung *et al*., 2017^4^. **(B)** Schematic of optogenetic targeting of the discrete PFC subsections: viral injection promotes opsin expression; blue light stimulation via optic fiber. **(C)** Histological slices (bottom) from injected and implanted PL, IL, and VO rats; eYFP-ChR2: green, DAPI: blue, scale bar: 1 mm. **(D)** 30-Hz blue-light stimulation effects in separate PFC subsections on long-delay trials over all sessions in each ChR2 cohort. Effects from previously published data^4^ for each corresponding subarea inhibited via continuous yellow-light illumination of NpHR-injected rats are plotted in lighter shades. Difference (laser vs. no-laser) in session-wise performance. Shape widths indicate session-wise distribution. Median value: black bar. (*) p <0.0083, (**) p <0.0017, (***) p <0.00017. See also Table S1. **(E)** Difference (laser vs. no-laser) in premature error rate under yellow-light control or blue-light stimulation at various frequencies for each PFC cohort. Black stars: significance compared to control within the same PFC cohort. Colored stars represent significance between different frequency parameters within the same color-coded cohort. (*) p <0.05, (**) p <0.01, (***) p <0.001. See also Table S2.

In detail, we explored the underlying information flow that might have led to the convergence of the excitatory effects among PFC subsections. To do this, we combined optogenetic modulation— facilitating reversible cell excitation for within-animal effect comparison—with behavioral and fMRI measurements to investigate the impact of PFC-subarea stimulation on motor control and evoked neuronal responses. Identifying potential network nodes underlying this effect via opto-fMRI, we revealed a common activation volume among the subsections, spanning the prefrontal cortex, olfactory bulb, basal forebrain, and secondary motor cortex. Notably, IL excitation led to the smallest but most precisely localized activation volume. At the same time, IL excitation produced the most robust behavioral effect (quantified via error rates and reaction times as behavioral indicators of motor inhibition in rats performing a response-preparation task). We further investigated functional synchrony within this region through multisite in-vivo recordings, uncovering performance-specific increases in PL-IL delta phase locking. Principal component analysis of single-unit activity during behavior further highlighted PL to have the most homogenous input-driven activity. We summarize our findings in a simple model of PFC subsections that suggests PL as input-dependent switch between motor inhibition and execution, modulated by an IL-dominated network supporting goal-directed action while keeping the behavior flexible.

## Results

### Excitation of individual PFC subsections induces a reduction of premature responses during long-delay trials

Goal-directed action control requires the inhibition of a response until the appropriate moment. To investigate this ability, we trained rats in a response-preparation task involving short- and long-delay periods. Long delays (targeting proactive inhibition) afford more time and information for preparation, leading to shorter reaction times, compared to short delays (targeting reactive inhibition)^4,7,8^. We then investigated the effects of PFC optogenetic manipulation on behavior, quantifying changes in error rate and reaction time to assess task performance (Figure 1A-B). We have previously demonstrated distinct behavioral effects in this task through subsection-specific inhibition of the PFC using Halorhodopsin (NpHR)^4^; prelimbic (PL) and infralimbic (IL) cortex inhibition revealed opposing effects on long-delay trials, while that of the ventro-orbital (VO) cortex influenced performance in short trials. To evaluate the potential opposing effects of excitation compared to inhibition, we used excitatory Channelrhodopsin 2 (ChR2; Figure 1C-D). Comparing within-session non-stimulated and stimulated trials, activation of each area led to reduced premature responses during long-delay trials (Figure 1D; Table S1). IL-activation produced a median decrease of 79.99% in premature releases (p<0.0083). PL and VO activation also reduced this error type, with median decreases of 22.30% (p<0.0017) and 27.57% (p<0.00017), respectively. The decrease of premature releases in long-delay trials upon PL excitation was complemented by an increase in correct releases (p<0.0017). The IL (p<0.0083) and VO (p<0.00017) excitation-induced decrease in premature responses coincided with increased correct and delayed response rates in long-delay trials. Effects on performance were accompanied by a reaction time (RT) increase after 1s laser stimulation for both IL (median 250 ms, p<0.005) and VO cohorts (median 80 ms, p<3.33e-04; Figure S1A, Table S3). Notably, IL-activation produced the largest effects on performance and RT. There were no robust effects observed in performance during short-delay trials (Figure S1B; Table S1). These findings suggest that PFC-ChR2 stimulation recruits a circuitry primarily involved in the proactive task component. Comparing our behavioral effects with previously obtained behavioral data from inhibition experiments^4^ (Figure 1D, S1B), only PL modulation showed opposing ChR2 *versus* NpHR effects.

### Higher stimulation frequencies induce larger behavioral effects and activation volume

Unlike inhibition, optogenetic excitation led to analogous behavioral output across all three PFC subsections. Previous studies have established ChR2 frequency-dependent neuronal response, activation patterns, and behavioral effects^9–11^. We confirmed frequency-dependence of the behavioral effect, suggesting efficient ChR2 targeting of a shared hub that integrates inputs from the PL, IL, and VO, resulting in similar excitation effects from each subarea. Control measurements revealed no significant effects (Table S2, Figure S1C). Intra-cohort variations between the control and all blue-light parameters for the most robust performance effect—a decrease of premature releases during long-delay trials—increased with higher stimulation frequencies (Figure 1E; Table S2).

The investigated PFC subsections share several functional downstream targets^12–14^, all of which represent potential network convergence points. To identify regions of interest underlying the analogous excitatory behavioral effects, we mapped whole-brain activation patterns during ChR2 stimulation using functional magnetic resonance imaging (fMRI; Figure 2A). This allowed us to characterize PFC subgroup-recruited whole-brain activity by measuring blood oxygen level-dependent (BOLD) contrast, reflecting neuronal activity^15,16^. We first validated ChR2 frequency-dependent activation in this experimental context (Figure 2B, Figure S2), observing a trend across all cohorts (n.s.) from 10–30 Hz for increased total volume activation, more recruited areas, and higher ipsilateral activation, with contralateral activation at higher frequencies. Notably, the PL showed a trend for higher activation volumes, and elicited cingulate cortex activation at all frequency parameters (Figure S2A), supporting previous findings of PL-ACC connectivity^1,17^ relating to action control.

**Figure 2:**
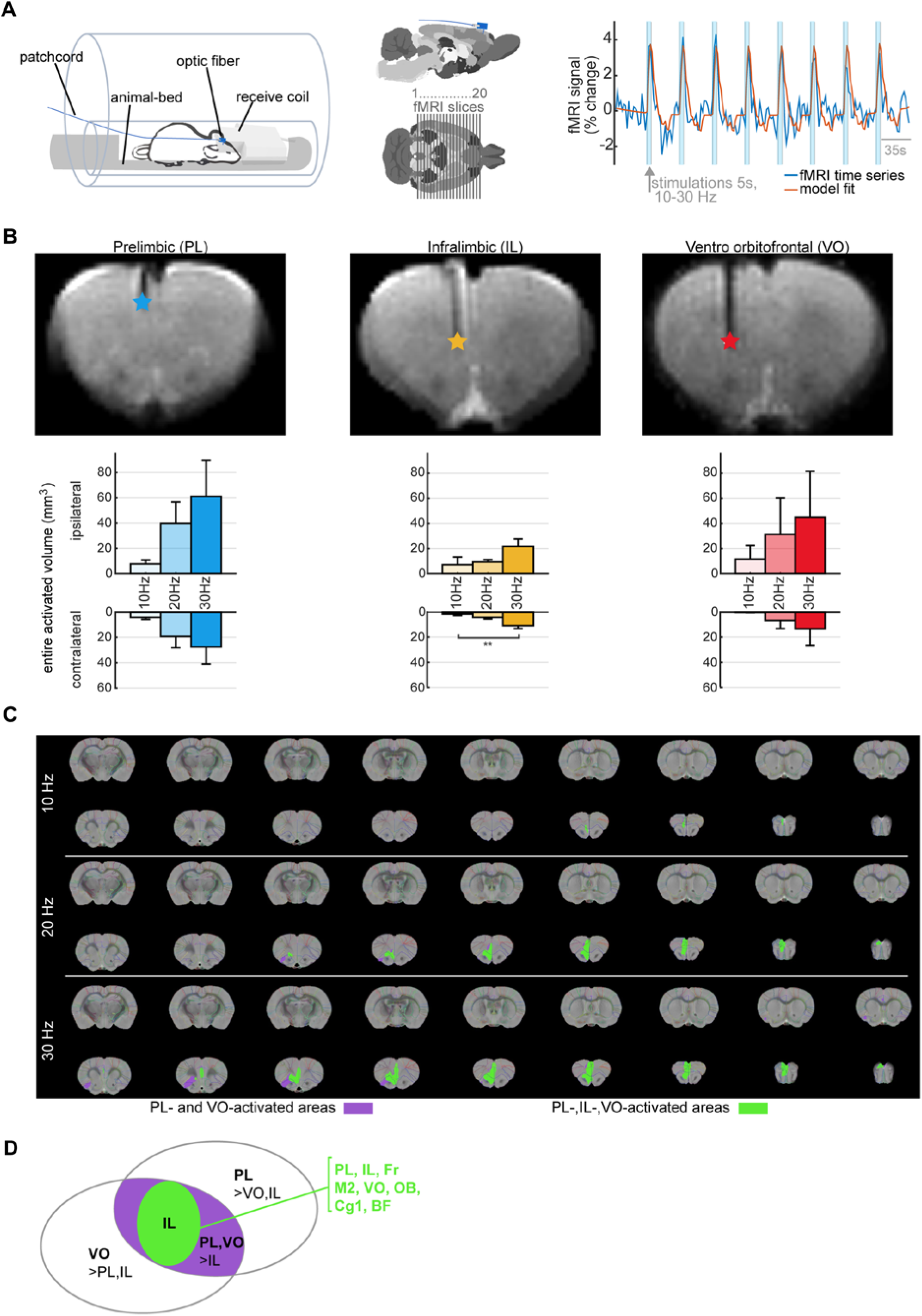
Opto-fMRI activation volume. **(A)** Opto-fMRI setup (left); schematic of low-profile fiber and patchcordrelative to the brain and fMRI slice positions (middle). Stimulation parameters for ChR2-injected rats during fMRI with BOLD fMRI activation marked in the time series, and evaluated via general linear modeling (right). **(B)** MRI images (top) of coronal slice with fiber and target stimulation area. Optical fiber tip identified by a colored star targeting individual PFC subareas. Respective whole-brain activation volume (mean ±SEM) of significantly activated voxels in the hemisphere ipsi- and contralateral to the optic stimulation site for 10–30 Hz stimulation, from group-level activation maps for PL, IL, and VO (bottom); see also Figure S2. Blue: PL, yellow: IL, red: VO. **(C)** Areas activated by multiple stimulation cohorts. Purple: shared PL and VO stimulation. Green: shared PL, IL, and VO stimulation. **(D)** Graphical summary of fMRI activation areas. ‘>’ indicates segregated activation areas with significantly stronger signal intensity in one group.

### Optogenetically-induced fMRI patterns indicate a common recruitment of IL-activated areas irrespective of stimulation location

In light of analogous ChR2-evoked behavioral effects, we compared group-level fMRI activation maps (S2C) to identify potential convergence points of ChR2 stimulation in the PL, IL, and VO (Figure 2C-D, S3A-J). At 10 Hz, activation patterns had only minimal overlap in the prelimbic system (spanning PL and IL) and anteromedial frontal association cortex (Fr), secondary motor cortex (M2), and olfactory bulb (OB). However, at 20-30 Hz, a larger common activation volume emerged, encompassing PFC fiber-tip-implanted regions and extending into the primary cingulate cortex (Cg1), OB, basal forebrain region (BF), Fr, and M2. The IL-recruited volume was precisely confined within the overlapping PL and VO activation areas and did not activate any areas more prominently than PL or VO stimulation (Figure S3A-F). Small volumes shared only across VO and PL stimulation were also identified, spanning the ventrolateral PFC, OB, secondary cingulate cortex (Cg2), and entorhinal cortex (Ent) (Figure S3G-J). Control experiments with eYFP-injected (ChR2-negative) and PFC-fiber-implanted rats showed no fMRI activation near the fiber tip, ruling out blue light heating effects (Figure S3K-L). In summary, IL-stimulation-activated areas were also activated by PL and VO stimulation, with the volume expanding at higher stimulation frequencies.

### Multisite recordings reveal performance-specific PL-IL oscillation synchrony

We aimed to characterize behavior-specific interregional communication within the opto-fMRI-identified regions of interest by analyzing oscillation synchrony, which facilitates cognitive processes such as information integration, attention, and action control^17–20^. Oscillatory activity within the mPFC has been implicated in discrete behaviors, including sensory-response-linked delta activity and waiting-specific theta activity^21,22^. Although not yet identified between the PL and IL, heightened delta phase synchrony has been identified within the PFC (PL-ACC) for appropriate responses^17^. Among all the PFC subareas tested, only the PL and IL affected proactive stopping behavior during both excitatory and inhibitory modulation. Given these robust effects, along with documented interconnectivity^23–26^ and interdependent oscillations^27^ between these two areas, we hypothesized that PL-IL synchrony in delta and theta bands may be necessary for task performance.

Using multisite PL-IL electrophysiological recordings during behavior, we specifically investigated synchrony during the modulation time window from the optogenetic-behavioral experiments. This allowed us to assess specific time points within the perturbation window from the opto-behavior experiment that are necessary for appropriate responses (Figure 3A-B). This approach revealed modulation of PL-IL delta phase-locking values (PLV), with a higher PLV at correct release (median: 0.7123, p=0.0431, z-value 2.0226, Figure 3C-D), compared to correct press (median: 0.5106) or premature release (median: 0.5606). In premature long trials, delta synchrony persisted at a low level during the press, without a clear pre-release increase. We did not observe consistent modulation of theta synchrony aligned to specific task stimuli.

**Figure 3:**
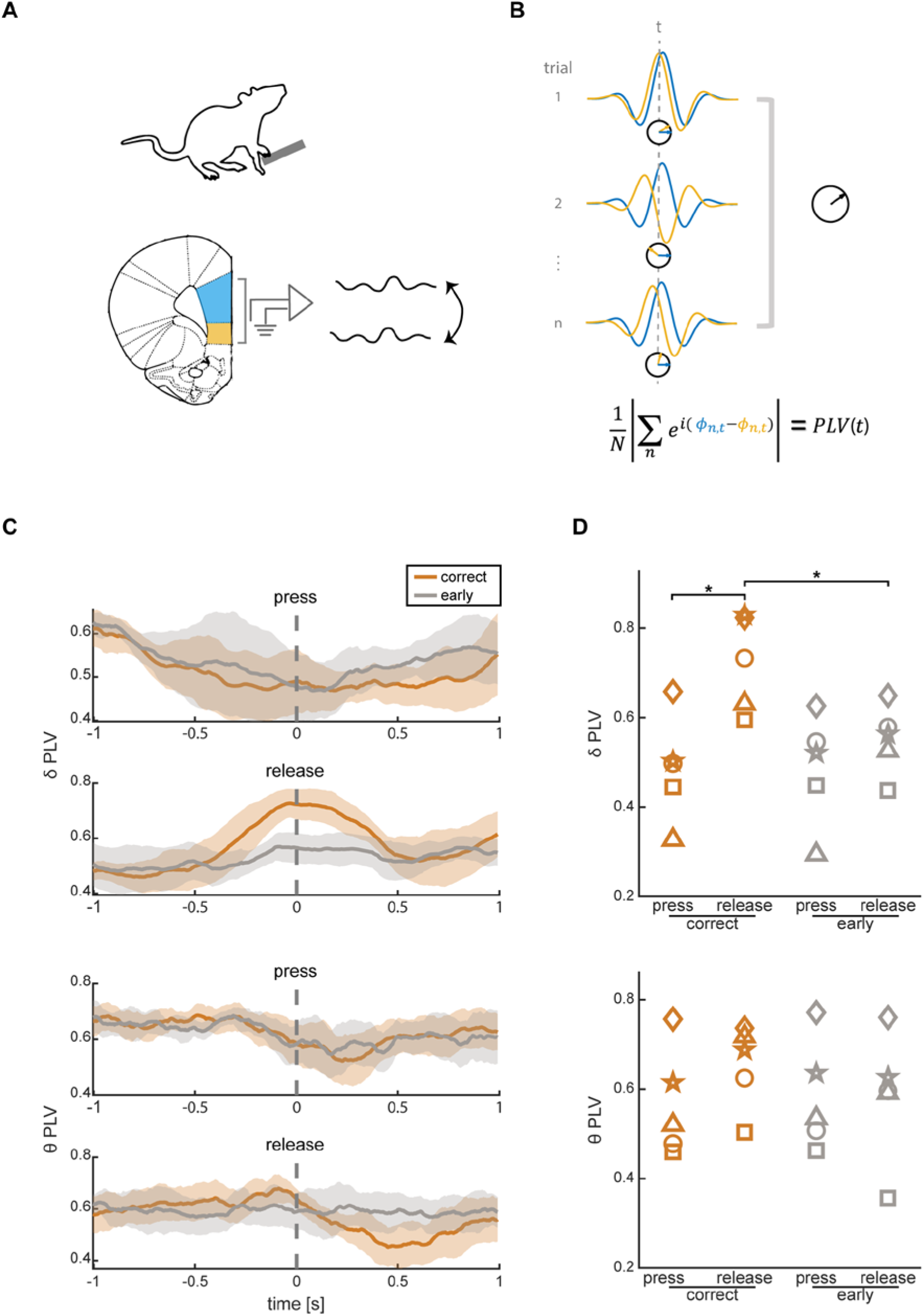
Performance-specific PL-IL synchrony. **(A)** Schematic of behavioral setup with multisite electrophysiological recording of PL and IL. **(B)** The schematic of the PLV estimation. Band-filtered LFP signals from a set of N trials (correct long or premature long) and two areas (blue and yellow) are aligned to press or release; PLV is computed for each time (t) around the alignment point (t=0). **(C)** Average PLV trace (dark line) ± SEM (shaded area) across animals (N=5) for correct (orange) and premature (grey) trials in delta (top) and theta (bottom) bands. **(D)** Average PLV at the time of press and release for delta (top) and theta (bottom) for individual rats (represented by different shapes) in same colors as in Figure 4C. (*) p<0.05

### Principal component analysis of neuronal activity during behavior indicates coherent input to the PL

While PL-IL synchrony offers a mechanism for the uniform behavioral effects across PFC subsections observed after ChR2 modulation, the discrete NpHR-evoked effects prompted further inquiry. Specifically, only PL inhibition displayed opposing behavioral effects to PL excitation, whereas perturbation of IL and VO did not result in a clear excitation-inhibition dichotomy. We hypothesized that the PL receives more temporally coherent inputs, allowing continued functional integrity during ChR2 activation, while NpHR modulation leads to disruptive effects. Conversely, the IL and VO likely receive more varied and heterogeneous inputs^4,29^; excitation therefore interferes with more diverse signals, leading to confounding effects. Notably, VO inhibition did not affect long-delay trials, suggesting a limited role in proactive task components. This potentially reflects greater neural variability in long-delay trials; we thus expected long-delay trial activity would be most variable in the VO, followed by the IL, with the PL the least variable. To confirm this, we analyzed neural recordings^4^ of individual subregions from rats performing in the response-preparation task (Figure 4A) using principal component analysis (PCA) to assess activity during correct long-delay trials. Explained variance per principal component (PC) was highest for PL, followed by IL, and then VO (Figure 4B, S4A-B), indicating that neural activity in the PL is more homogenous. We identified PCs 1-3, which had greater explained variance than the corresponding PC from randomized datasets (Figure S4B), as significant and retained these for further analysis. An error trial analysis of PC-transformed activity revealed performance-dependent neural activity (Figure 4C). To check if the significant PCs resulted from small but strong outlier populations or from a uniform population response, we analyzed how correlated each neuron was to the corresponding PC traces. Correlation of individual units over significant PCs (1-3) in each area revealed the PL to have the highest ratio of neurons with correlation magnitudes outside the 95% of neuron correlation magnitudes from shuffled data; this was followed by the IL and VO (Figure 4D). This indicates that a larger proportion of neurons in the PL are strongly aligned with these principal patterns of activity, further revealing the PL’s comparatively homogeneous activity of the three PFC subareas. Importantly, a coactivation analysis (Figure S4C) did not reveal any significant task correlation of ensemble activity in any PFC subarea, further suggesting that the homogeneity observed in the PL is primarily input-driven rather than a result of internally coordinated activity. This supports our model of distinct task-correlated variability of neural processing in PFC subsections, where neural and behavioral outcomes are predominantly shaped by a temporally coherent depolarization that is most likely due to afferent activity.

**Figure 4:**
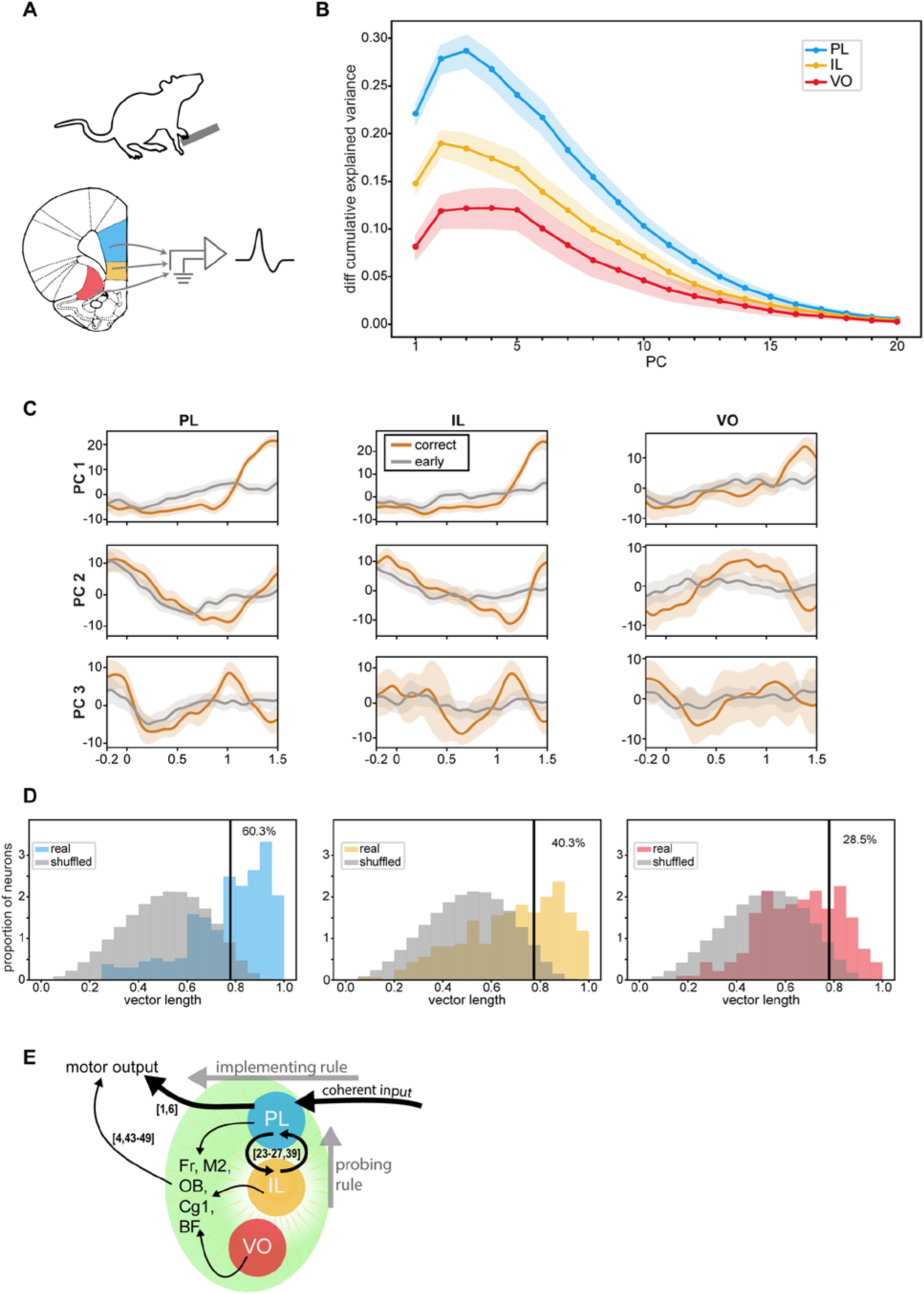
PFC subsection local dynamics reveal variable input patterns. **(A)** Schematic of behavioral setup with individual PFC subsection electrophysiological recording. Experimental data from Hardung et al^3^. **(B)** Difference of observed cumulative explained variance ratios and random cumulative explained variance ratios (mean difference between real explained variance and random explained variances from shuffled datasets, represented by the dark line; the 95^th^ percentile is represented by the lightly shaded region). PCA was performed on neural activity of long correct trials (see also Figure S4A-B). **(C)** For significant PCs 1-3 (top-bottom), PC-transformed correct (orange) vs error (grey) trial neural activity aligned to press (t=0) for each subsection, illustrating performance-specific neural activity. **(D)** Normalized neuron proportion for binned correlation circle vector lengths over significant PCs 1-3 (see Figure S4C) across each subsection, as well as for shuffled neural data (grey) from each subsection. Percentage of neurons with vector lengths that above the 95^th^ percentile (indicated by black bars) of lengths from the random data is indicated on the graph. **(E)** Schematic illustrating connectivity patterns revealed in our study. Black arrows indicate connectivity demonstrated by our experimental data or well-evidenced in previous studies (cited in brackets). PL, IL and VO activation recruits a local shared activation area (light green) including PL, IL, VO, Fr, M2, OB, Cg1, and BF; these regions are implicated in motor output^4,43–49^, and represent a putative target region linked to the shared behavioral output of the three PFC subareas. Within this region, PL-IL synchronized activity was identified for correct motor output in the task, in line with well-described PL-IL interconnectivity^23–27,39^. Coherent input drives the homogenous activity observed in PL during lever pressing, facilitating the PL’s direct control of motor output^1,6^. The IL and VO had more heterogeneous activity, driven either by less coherent input or by internally coordinated variability.

## Discussion

This study elucidates the bidirectional effects of optogenetic manipulation of PFC subsections on motor inhibition and specifically explores their network convergence. Opto-fMRI measurements highlight the overlapping activation of cortical regions, contained within the IL-activated volume. Meanwhile, analogous behavioral output occurred upon excitation, irrespective of the specific subsection targeted, suggesting network convergence within the PFC. Both the behavioral effect and fMRI volume increased at 20-30 Hz stimulation, indicating efficient ChR2 targeting of a shared hub. The overlapping activation volume among the three cohorts does not implicate shared downstream targets, but rather local cortical regions connected to PFC^13,30–32^. This indicates that PFC activation contributes to the integration of these regions within a broader cortical network. As a caveat, the use of anesthesia during fMRI may have limited the detection of downstream activation targets^33,34^ such as the striatum or thalamus. However, this setting still allows for the evaluation of cortical effects of the stimulations, as well as the side-by-side comparison of stimulation of different PFC subsections. Through this method, we found that IL stimulation uniquely recruited the common activation area, indicating its strong modulatory role in this circuit (Figure 3D).

Building on these findings, we focused on behaviorally relevant neural integration. Given the well-described PL-IL connectivity^23,24^ and the IL’s prominent recruitment of the shared activation area, we examined oscillation synchrony within this subset, identifying performance-specific PL-IL delta phase-locking modulation. This aligns with previous research linking delta synchrony within the mPFC to cue-driven responses^17,20,21^. fMRI data revealed robust PL recruitment of the cingulate cortex, implicated in cue-driven response and top-down control of the motor cortex^22^, further complementing our observed performance-specific delta synchrony prior to cue-driven responses.

We propose that PL-IL synchrony facilitates cognitive control; disrupting this synchrony likely impairs coordinated activity, leaving PFC subsections to independently regulate behavioral output. Investigating neural patterns of individual PFC subsections revealed that the PL is driven by coherent inputs, indicated by more homogenous neural activity (i.e., higher explained neuronal variance per PC and more neurons correlated to significant PCs) without task outcome-related ensemble activity. This suggests that the bidirectional effects of PL modulation arise from targeting uniform inputs with both NpHR and ChR2. These findings underscore that coherent input to the PL is crucial for its direct control of the motor system^1,4,6^, providing stable and consistent neural signals necessary for precise motor regulation.

Our findings highlight both intra-PFC network convergence and distinct roles for PFC subsections in cognitive control. Previous work has described the functional dichotomy between the PL and IL—also termed the dorso-ventral mPFC gradient—exerting opposing influences on cognitive function^35–38^. This bidirectional signaling extends to goal-directed action, reward seeking, fear learning, and pain modulation^35–38^. However, recent evidence emphasizes the role of intrinsic reciprocal PL-IL connectivity for adaptive behavior^23,26,39–41^. The shared ChR2 behavioral effects and activation volume observed alongside performance-specific synchrony within the PFC highlight the relevance of such cortical-cortical connectivity; the discrete NpHR behavioral effects and PCA results argue for the relevance of subsection-specific neural patterns. This underscores the importance of considering the individual contributions of PFC subsections in cognitive control and challenges the practice of treating these regions as a single mPFC entity in experimental design and interpretation. It also illustrates that opto-inhibition is crucial for discerning the network activity necessary for specific tasks, as it selectively disrupts relevant connectivity; excitation alone can obscure these specific contributions by non-selectively activating broad neural pathways.

Together, our results propose a new simple model of the mPFC (**Fig 4E**): We have provided evidence that the PL receives coherent input, which renders it a decisive “switch” between motor execution and inhibition. The PL is suited to this role due to its direct contribution to motor output^1,6^. A properly functioning (i.e. undisturbed) PL implements correctly timed actions. Simultaneously, the IL has a strong modulatory effect on the PL, working against learned rules that are implemented by the latter. If the IL influence on the network is disrupted, the PL works in isolation and thus implements correctly timed actions according to learned rules. We hypothesize that PL-IL interaction allows for flexible behavior^3^, as indicated in fear extinction studies^19,25,26,36,42^, for instance. To ensure that the IL does not block all reasonably learned rules, a fine balance between the PL and IL needs to be implemented. Our model suggests that this balance is derived from the degree of coherence of input into the PL: Under physiological conditions (i.e., without optogenetic disturbances) the more coherent the input to the PL is, the less ambiguous the situation becomes. In these cases, IL’s influence is limited. The less coherent the input to the PL (i.e., the more ambiguous the situation is), the stronger the “disruptive” IL network effect becomes.

Overall, our study demonstrates fine-tuned functional cortico-cortical synchrony in motor control, as well as discrete roles of PFC subsections. This lays the groundwork for future investigations into context-specific intra-cortical activity and reciprocal PL-IL connectivity.

## Methods

### Ethical Statement

All animal-related procedures adhered to guideline RL 2010 63 EU and were approved by the Regierungspräsidium Freiburg (TVA G-20-26, TVA G-20-65). All rats were maintained in a reversed 12-hour light cycle, with free access to food and water. Rats involved in behavioral experiments underwent water deprivation during behavioral training.

### Behavioral Training

Ten rats (male Sprague Dawley, 6 weeks old; Charles River) were familiarized with the response-preparation task in chambers containing two retractable levers, a house light, two cue lights, and a reward port connected to a variable infusion rate syringe pump (Med Associates Inc., USA; Figure 1A). Familiarization involved adjusting reward amounts, tone delays, and randomization of the tone delays to sustain motivation until rats successfully performed under the final task parameters. The session start was signaled by house light activation and simultaneous release of both levers into the chamber. Cue lights above each lever indicated the active lever, alternating between trials. Rats pressed a lever (50 g tension) until a tone (3.0 kHz, 80 dBA, 0.5 s) occurred. Lever release within 0.5 s of the tone resulted in reward delivery (Figure 1B), activating a syringe pump (0.05 ml/s for 0.5 s) and delivering 0.025 ml of 3–6% sucrose solution into the reward port. Initiating a new trial required head entry into the reward port, activating a new trial via a movement sensor. The delay between the lever press and the tone was either 0.3 s (representing the reactive element of the response task) or 1.0 s (representing the proactive element). These delays were pseudorandomized in intermixed blocks of a higher probability of long delays (75% long delays and 25% short delays) or an equal probability of long and short delays (50% long delays and 50% short delays). We avoided using blocks with a high probability of short delays, as it led to the rats developing bias for short-delay trials. The blocks switched after 15 correct trials. Incorrect trials incurred a “time-out” of 1.0 s, during which head entry to the reward port did not start a new trial. Each behavioral session lasted 30–60 minutes, depending on the rats’ performance and motivation. After reaching a minimum performance of 55%, they underwent stereotactic surgery for viral injection, fiber implantation, and behavioral recordings during optogenetic stimulation. A subsequent cohort of five rats (female Sprague Dawley, 20–25 weeks old; Janvier Labs) underwent training in the established protocol using a modified experimental setup featuring only one lever, implemented to expedite the training protocol; this cohort underwent silicone electrode implantation for behavioral recordings alongside electrophysiological measurements.

### Stereotactic Surgery

All rats were briefly anesthetized with 5% isoflurane (using oxygen (O^2^) as a carrier gas), followed by intraperitoneal (i.p.) injections of 80 mg/kg ketamine (Serumwerk Bernburg, Germany) and 0.06 mg/kg medetomidine (Dextomidor, Zoetis Deutschland GmbH, Germany). Subcutaneous (s.c.) 0.05 mg/kg buprenorphine (CP-Pharma, Burgdorf, Germany) was administered pre-surgery. During surgery, anesthesia was maintained at 0–3% isoflurane and 0.5 l/min O^2^, rats were kept on a heating pad, and their temperature was monitored via a rectal sensor. Eye moisture was maintained using ophthalmic ointment (Bepanthen, Bayer Health Care, Germany). We applied 1– 5 ml of saline s.c. injections (isotonic NaCL 0.9%, B. Braun Melsungen AG, Germany) every two hours to maintain fluid balance. Two ml of 5% glucose solution (B. Braun Melsungen AG) was injected s.c. after 4 hours of intervention. Fur was shaved at the surgical site, and the rats were then head-fixed into a stereotaxic frame (World Precision Instruments, Germany). The surgical site was disinfected using Kodan (Schülke, Germany) and Braunol (B. Braun Melsungen AG, Germany). Lidocaine (2% xylocaine gel, Aspen Pharma Trading Limited, Ireland) was applied to the incision site (maximum dose 10 mg/kg) 5 minutes prior to making a maximum 2 cm-long incision along the skull, from anterior to posterior. The edges of the skin were held apart with surgical clamps. Subcutaneous tissue was removed from the skull with a bone scraper, followed by the removal of leftover debris or tissue with 3% hydrogen peroxide. The head was leveled in the anterior-posterior (AP) direction to ensure bregma and lambda were within a 0.05 mm height difference of each other. Medial-lateral (ML) leveling was achieved by positioning the head so that ML locations to the right and left of bregma were within 0.05 mm of each other in the AP plane. A craniotomy was drilled around the target area of injection or implantation.

The behavioral cohort (male Sprague Dawley rats, 14–18 weeks old, 400–620 g at surgery; Charles River) underwent bilateral viral injection and bilateral fiber implantation. After craniotomy, a glass microcapillary (tip diameter ≃35 μm; 1–5 μL Hirschmann® microcapillary pipette, Z611239, Sigma-Aldrich, USA) attached to a stereotactic holder was lowered to the injection site. A total of 300–400 nL of a viral vector (pAAVhSyn-ChR2 (H134R)-eYFP, pAAV-hSyn-NpHR-eYFP, or pAAV-hSyn-eYFP; University of North Carolina Vector Core, USA) was injected via a custom Openspritzer pressure system^50^. The rats underwent subsequent bilateral implantation with 200 µm-wide fibers fixed into 230 µm-diameter stainless steel ferrules (Precision Fiber Products Inc., USA). To stabilize the implants, we added miniature self-tapping screws (J.I. Morris Company, USA) to the skull. A fiber was inserted with a stereotactic holder through the craniotomy into the desired implantation area; the craniotomy was then filled with silicone elastomer filling (Kwik-Cast, WPI, Sarasota, FL, USA) to prevent brain swelling. We then applied a thin layer of Super Bond C&B cement (Sun Medical Co., LTD, Moriyama City, Shiga, Japan) around the fiber for initial stabilization and formation of a base on the skull. Paladur (Heraeus, Hanau, Germany) was then layered to create a sturdy skull implant. The surgical site was sealed around the implant with medical sutures (DS16 Dafilon, B. Braun, Germany).

The multisite recording cohort (female Sprague-Dawley rats, 20-25 weeks old, 300–340g at surgery; Janvier-Labs) underwent silicone electrode implantation (E32+R-100-S2-L6-200 NT, Atlas Neuroengineering, Belgium) for simultaneous electrophysiological recording in the PL and IL during performance in the response-preparation task. The implantation protocol was identical to the above, except without viral injection and with the implantation of electrodes instead of fiber. Additionally, one miniature self-tapping screw was inserted slightly lower to penetrate the cortex; this screw was attached to a ground wire that was later fully embedded within the implant cement.

fMRI experiment rats (male Sprague Dawley rats, 7–8 weeks old, 288–450 g at surgery; Charles River) underwent unilateral viral injection and implantation under a similar protocol as the behavioral experiment animals, with some differences. We did not use metal screws as an anchor for the implant; instead, the skull surface was roughened with a skull scraper to allow the Super Bond cement to efficiently attach to the surface. Additionally, a low-profile fiber stereotactic holder (Doric, Canada) was used to insert one (unilateral) low-profile fiber construct consisting of a 200 µm-wide fiber attached to a 1.5 mm-diameter zirconia ferrule (Doric, Canada) to maximize proximity of the MRI coil to the rat skull during experiments (Figure 1F and 1G).

### Post-Operative Care

An s.c. injection of carprofen (5 mg/kg) was applied immediately after surgery, followed by additional doses 1–2 times daily for 1–3 days post-surgery (partially substituted by oral doses). Additionally, wet food was given 1–2 times daily to all rats for up to 1.5 weeks post-surgery to ensure adequate nutrition and facilitate recovery.

### Stimulation and Behavioral Recording

Following the post-surgical recovery period, rats were retrained for 1–2 weeks before proceeding to behavioral recording sessions with optogenetic stimulation. Three of the operated rats were excluded from retraining and behavioral recordings (two experienced surgery-related complications and one had an unstable implant post-surgery). This left three PL, three IL, and four VO rats for behavioral recording. We performed behavioral recordings under optogenetic stimulation for 10–20% of the trials per session. During laser-stimulated trials, the laser was applied from the time of the press until the release or the tone arrival, whichever occurred first. Sessions lasted 30–60 minutes, depending on behavior and performance. The fibers in the implants were coupled to a LightHUB compact laser combiner (OMICRON Laserage, Germany) via two mono fiberoptic patchcords and a fiber-optic rotary joint (FRJ_1x2i_FC-2FC, Doric Lenses, Canada). Optogenetic excitation was achieved with blue laser light (λ = 473 nm, 5–8 mW, measured at the tip of a 200 µm-diameter fiber with a PM100D power meter; Thorlabs, USA). Stimulation frequency was constant over each session and varied between sessions (2 ms-wide pulses at 10, 20, or 30 Hz). Control parameters were applied by yellow-light stimulation (λ = 594 nm, 10–14 mW), which should not activate ChR2, to test for any confounding effects of heating or light application during the task. We tested different yellow-light parameters, to ensure that pulsed vs continuous light did not create any effects on behavior (Figure 2B). We used 20 Hz yellow light (2 ms pulses) for PL rats; control parameters for the IL group were 30 Hz yellow light (2 ms pulses); VO rats were stimulated with continuous yellow light for control measurements. NpHR-implanted rats from previously published work in our group^4^ also underwent blue-light controls (λ = 473 nm, 10-14 mW), as this is non-activating for NpHR (Figure S1C). Sessions were excluded from future analysis if the patchcord output was compromised, indicated by loosening of the patchcord-ferrule connection, or damage to the patchcord during the course of the session (monitored by light output measurements before and after each recording session).

### Multisite Electrophysiological and Behavioral Recording

Electrophysiological data was acquired via RZ2 Bioamp (Tucker-Davis technologies, TDT) hardware and the dedicated OpenEx Software Suite. The implanted electrode-EIB assembly was connected to the recording equipment through the ZD32 ZIF-Clip® digital headstage. The signal was sampled at the rate of 24,414Hz, filtered online and the down-sampled (976.56Hz) LFP component (0.1–300Hz) saved as a separate data stream. In addition to the TDT setup, control of behavioral variables and equipment was aided by Med-PC (Med-Associates Inc.) and custom-written MedState Notation programs. Digital streams between TDT and Med-PC allowed for event synchronization and offline analysis.

### Perfusion and Histology

The behavioral experiment rats were euthanized via an i.p. injection of 400 mg/kg sodium pentobarbital (Release®500, WDT, Germany), while fMRI rats were euthanized via i.p. injection of ketamine and xylazine (due to regulation/animal protocol changes in the time between these experiments). Both cohorts underwent transcardial perfusion with PBS, followed by ice-cold 4% paraformaldehyde. Following perfusion, the brains were removed and post-fixed for 1–2 days before transfer to 30% sucrose (Merck KGaA, Darmstadt, Germany) dissolved in PBS, and stored at 4°C. Once fully saturated with sucrose, the brains were sectioned into 50 µm slices using a sliding microtome (Leica model SM2010 R, Leica Biosystems, Germany). The slices were then mounted on coverslips with Vectashield Mounting Medium including 4′,6-diamidino-2-phenylindole (DAPI, Vector Laboratories Inc., USA). Imaging was conducted using an Axioplan 2 Imaging fluorescence microscope (Zeiss, Germany).

### Behavioral Analysis

Behavioral recordings were analyzed with custom-written MATLAB (MathWorks, USA) scripts. Although rewarded as correct trials during behavioral sessions, trials with a reaction time (RT) shorter than 150 ms were considered premature releases; due to auditory processing delays^2^, it is assumed that the rats could not actively respond to the tone this quickly.

The behavioral data from stimulation sessions were analyzed to assess changes in error rate (Figure 1D-E, Table S1). Percent error rate was calculated as the percentage of error types presented across all trials of one type, for example, delayed responses in unstimulated long-delay trials. The difference in error rates between stimulated and non-stimulated trials was calculated for each trial outcome type in single sessions, and the median percent change in each trial outcome was calculated across all sessions for each group of rats according to the targeted PFC area. A Wilcoxon signed-rank test was employed to check for differences between stimulated and non-stimulated error rates across all sessions in each cohort, with Bonferroni-adjusted p-thresholds (Figures 1D, Table S1). One-way analysis of variance (ANOVA) with subsequent Tukey-Kramer multiple comparisons was used to test for the variance of laser effects on error rates between control and various frequency stimulation parameters within each cohort (Figure 1E Table S2). RT was also investigated (Figure S1A, Table S3); this was defined as the time from the tone until lever release, pooled for trials that had a tone (both correct and late). For each session, the difference of the mean RT for short- or long-delay trials between laser-on and laser-off was calculated; the median of the RT difference was calculated for each stimulation parameter over all sessions within each cohort. Significance between laser- and non-laser-stimulated RTs for the same trial type was tested with a Wilcoxon signed-rank test and Bonferroni-adjusted p-thresholds. Violin plots were visualized with custom-written MATLAB scripts^51^.

### Synchrony analysis

Simultaneous recordings from PL and IL during the execution of the behavioral task were analyzed for LFP synchrony. Consistent phase differences over trials were captured using phase-locking value (PLV) analysis, which assesses synchrony by measuring the mean phase difference between PL and IL signals across trials (Figure 3B). The magnitude of the mean vector represents the strength of synchrony, ranging from 0 to 1, with higher values indicating consistent phase differences across repeated trials at the time points of interest. Analysis was conducted separately for each animal, alignment (press/release), and outcome (premature/correct). Trials from sessions across different days (4–5 per animal) were combined for statistical power, with correct trials and premature trials with a minimum 300 ms holding time pooled. Identical electrode pairs (the most distal electrodes on the probe (1.4–1.5 mm apart, one located in PL and one in IL) were chosen across these different sessions for each animal to ensure sampling from identical populations.

All analyses were performed with custom MATLAB scripts and the Fieldtrip toolbox^52^. LFP data from the electrode pairs were first down-sampled to 244.14Hz, curated for large amplitude artifacts, and decomposed into a time-frequency representation using the Fieldtrip function ft_freqanalysis and the Hilbert method for specified frequency bands (two-pass FIR filters of orders 366 and 183 with passbands 2–4Hz (delta) and 4–8Hz (theta), respectively). The complex coefficients of both time series were then normalized to unity.

Given an alignment point *a∈A, A* = {*press*, *release*}, outcome *o∈O, O* = {*earlylong*, *correctlong*}, animal *r*, frequency band *f*, and letting *N* be the set of trials with outcome *o* we calculated the phase locking value for each animal, outcome, alignment, and frequency band as:

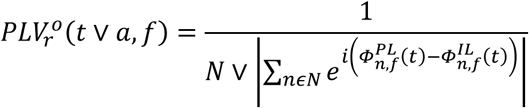

Where *t* is the timewith respect to the alignment point *a, N V* is the trial count of N, and *ϕ_n,f_* (*t*) are the phases of the spectral coefficients of each area on trial n.

To generate PLV synchrony spectrograms (Figure S4D), the LFP time series were decomposed with complex Morlet wavelets (width of 5 cycles, resolution of 0.5Hz), and the PLV calculated analogously. Although we employed PLV analysis for its comprehensive assessment of phase synchronization consistency at fixed time points across trials, we conducted a complementary analysis of our observations using the phase-locking index (PLI) to ensure the robustness of our findings (Figure S4D). The data for PLI spectrograms were decomposed in an identical manner as for the PLV analysis, and the PLI estimate was obtained using the following formula:

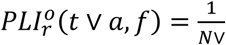

Where 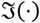 takes the imaginary component of the complex argument and *sgn*(·) the sign (±1) of its argument.

While comparing example data, we noted that higher PLV values were generally reflected by higher PLI values, albeit with a diminished effect. Lower PLI values are expected given PLI’s susceptibility to phase cancellation and its focus on the asymmetry rather than the consistency of phase differences. This cross-validation confirmed similar increases in the low-frequency bands, which were then extracted for statistical analysis.

Statistical analysis involved comparing PLVs at alignment points across 5 animals using the Wilcoxon signed-rank test. PLV was tested for press-to-release on correct trials, and release-to-release between correct and premature trials.

### Principal component analysis (PCA)

#### Data preprocessing

To analyze neural activity during the response-preparation task, we processed datasets acquired in a prior study^4^ of electrophysiological recordings in the PL, IL, or VO during the response-preparation task. Neurons with firing rates above 100 Hz or below 1 Hz were excluded. Correct long-tone delay trials were selected for further analysis. Spike data were binned into 10 ms intervals defined time windows ranging from 200 ms before to 1500 ms after the press for each trial were extracted, and then smoothed by a Gaussian filter with a kernel size of 50 ms. Peristimulus time histogram (PSTH) plots were then computed for each neuron over the time window. This resulted in a matrix with dimensions of (neuron count) × 170 bins, encapsulating the firing patterns of neurons within the specified time frame. This matrix of PSTHs was normalized by z-transformation and then processed via PCA to extract the PCs representing the major sources of variance in the neural activity dataset.

#### Bootstrap analysis for correct vs error activity comparison

The PSTH of all neurons (during long correct trials) within each brain area was computed as described above, from which all neurons recorded in that area were randomly drawn with replacement. This was repeated 10000 times to bootstrap samples. PCA was then conducted on each bootstrap sample. For comparison of the real data (correct vs error), the PCs, fitted on correct responses, were used to transform the correct and error response activity. This allowed visualization of the PC-transformed correct and error response averaged over the 10000 bootstrap iterations, to qualitatively assess if each PC is performance-related (Figure 4C). PCA maximizes variance without distinguishing between positive and negative directions, potentially yielding mirrored traces. To ensure consistency, bootstrapped traces from each iteration were correlated to a reference trace (the trace from the first bootstrap). Negatively correlated traces were flipped before averaging. The mean and 95^th^ percentile were then calculated over correct or error traces for each PC, providing estimates of the data distribution.

#### Shuffling procedure for comparison

A shuffling procedure was implemented to compare real and randomized PCs. Time bins throughout the entire session (overall firing rate for the whole session) for each neuron were independently shuffled, preserving the overall firing characteristics of each neuron while disrupting the temporal structure. This process was repeated 10000 times to generate randomized datasets. Mean firing rates were computed for each shuffling iteration, resulting in 10000 shuffled datasets (neurons × bins) per brain area. These datasets were then processed in the same manner as the real data (see above section: data preprocessing). PCA was performed on each of these smoothed and normalized shuffled datasets, deriving 10000 PCAs from shuffled data per brain area. The mean explained variance ratio per PC 95^th^ percentile was then calculated from the shuffled dataset, enabling comparison of the observed and shuffled distributions (Figures S4A), and providing insight into the presence of structured neural activity patterns beyond what would be expected by random chance. We additionally subtracted the shuffled explained variance of each of the 10000 iterations from the real explained variance per PC, to visualize the differences in explained variance per PC across PFC subsections (Figure 4B, Figure S4B). To identify significant PCs, we determined the components that had larger explained variance than their corresponding component from the shuffled data^53^ (Figure S4B).

#### Correlation analysis

To assess the relationship between individual neuronal activity and significant PCs, we conducted a correlation analysis^54,55^ via Pearson correlation coefficients. Specifically, we calculated the correlation between firing patterns of each neuron and the first three PCs within each brain area. For each neuron, the correlation coefficients were used to generate vector magnitudes within a correlation circle. The magnitudes of these vectors were binned into intervals (0.05) ranging from 0 to 1. This process was performed for both real and shuffled data (10,000 iterations). The raw binned neuron counts were normalized to account for the shuffle dataset iterations and different amounts of total neurons recorded from in each brain area. The distribution of vector magnitudes from the real data was compared to that of the shuffled data. We identified significance thresholds by calculating the 95^th^ percentile of vector lengths from the shuffled data. Finally, we determined the ratio of real neuron vector lengths that exceeded this 95^th^ percentile threshold, identifying neurons whose activity was strongly correlated with the significant PCs, revealing the degree to which neuronal activity in each brain area is associated with these PCs.

#### Activation Analyses

To identify neuronal assemblies during lever pressing in a behavioral task, we conducted an activation analysis^56^ of simultaneously recorded neural activity. Population vectors were constructed from this spike data for 50 ms time bins to represent the activity of all neurons at that specific time point. Each neuron’s activity was z-scored based on its mean and standard deviation during the entire session. We then performed a dimensionality reduction by diagonalizing the covariance matrix of the z-scored population vectors *p*(*t*). Eigenvectors *ν_i_* were retained only if their corresponding eigenvalues *λ_i_* exceeded the Marchenko-Pastur bound: 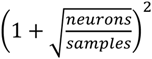. Activation strength *R*(*t*) for a population vector *p*(*t*) at time *t* was computed as *R*(*t*) = *p*(*t*)*^T^ Cp*(*t*), where C is defined as 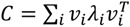. Significance of computed activation strengths was determined by comparison of *R*(*t*) values to the shuffled control. The shuffled control was produced by shuffling the time bins during lever pressing independently for each neuron 100 times. This allowed us to identify and quantify the activation of neuronal assemblies over time during the trial.

### Optogenetic MRI experiments

#### Animal Preparation and Maintenance

Rats were anesthetized with 4% isoflurane for ca. 90 s and then maintained at 1–2% isoflurane during preparation for the MRI measurement. The rats were secured on an animal bed, and a bolus of 0.05 mg/kg medetomidine (Domitor, Vetoquinol, Germany) was injected s.c. A catheter (outer diameter 0.6 mm) was inserted into the dorsal subcutaneous tissue of the rat to continuously administer medetomidine (0.1 mg/kg/hr) throughout the measurement. Isoflurane was discontinued at the beginning of the scan. Throughout the procedure, rats were supplied with air (30% O₂), with continuous monitoring of respiration and maintenance of body temperature. The total time of the MRI measurements was limited to 1.5 hrs. In the event of detection of movement during the scan, isoflurane was reintroduced, and the scan was halted. Post-measurement, animals received 0.1 mg/kg atipamezole (Antisedan, Vetoquinol, Germany) s.c. to reverse the medetomidine anesthesia and observed for 1 hour before being returned to the animal facility.

#### MRI Acquisition

MRI data were acquired on a preclinical 9.4 T system (BioSpec, Bruker BioSpin, Germany) with a volume transmit (diameter 7.2 cm) and surface receive coil (Bruker BioSpin, Germany). Anatomical reference images were acquired using a T2-RARE sequence with the following parameters: matrix size of 128 × 80, in-plane resolution of 0.25 × 0.25 mm^2^, 20 slices with a thickness of 0.8 mm, effective echo time (TE) 40 ms, repetition time (TR) 2.5 s. Data were acquired with a multi-echo (ME) gradient echo planar imaging (EPI) sequence with the following parameters: matrix of 64 × 40, in-plane resolution of 0.5 × 0.5 mm^2^, 20 slices with a thickness of 0.8 mm, TR of 2 s, and three echoes at TEs of 12 ms, 21 ms and 30 ms.

#### Optogenetic stimulation during fMRI

Stimulation parameters were configured with a Prizmatix pulser (Prizmatix Ltd., Israel), which controlled the LightHUB compact laser combiner (Omicron-Laserage Laserprodukte GmbH, Germany). Trigger-out signals from the MRI scanner during the fMRI scans were detected via a data acquisition board (DAQ, NI USB 6251 BNC, National Instruments, USA) using a custom-written MATLAB script. The DAQ transferred the trigger signal to the Prizmatix pulser to initiate laser stimulation.

A single-block design fMRI experiment comprised eight 5 s stimulation blocks and 35 s rest periods, with each measurement lasting 6 min. Over several measurements, we tested blue light stimulation in ChR2-expressing rats at various frequencies (10, 20, and 30 Hz) with 2 ms pulse widths. The light intensity was 12–13 mW at the fiber tip. Higher intensities (32–39 mW) were required for two animals to elicit activation; this was attributed to degraded ferrule flatness post-implantation, as light leakage was observed at the ferrule-patchcord connection.

In total, we measured 15 ChR2-injected rats. Five were excluded because we were not able to induce an fMRI activation with light, either due to a broken optical fiber, or misalignment of the fiber and viral expressing target area (identified during histology). Ten ChR2-injected rats (three PL-, four IL-, and three VO-targeted) were included for analysis. The blue light stimulation parameters were also investigated in YFP control rats, which received only YFP-expressing virus without opsin. Four YFP control rats (n = 4; two rats with fibers in PL, one IL, and one VO) were measured to assess potential confounding factors such as light-induced heating (Figure S3K-L)

#### MRI Analysis

The fMRI data were processed using FSL (FMRIB Software Library, v6.0, Oxford, UK) and custom-written MATLAB scripts (MathWorks, USA). Preprocessing included motion correction (using the FSL function “mcflirt”), slice-timing, and high-pass temporal filtering (100 s cut-off). Subsequently, the three individual time series (each for a unique echo time) were combined into a single “multi-echo” time series using a T2*-weighted summation of echoes^57^. The required T2* maps were calculated from an exponential fit of the mean signal of the first 20 volumes of each time series. Finally, the brain was masked, and spatial smoothing (1.5 times in-plane resolution) was applied.

To evaluate fMRI activation, first-level analysis was performed using FSL^58^, which evaluates each data set separately. Using FSL, the experimental design containing the timing and duration of stimulation and rest periods was convolved with finite-impulse-response (FIR) basis functions (8th order spread over a 24 s window), and the individual activation was assessed using general linear modeling (GLM)^59^. Subsequently, these individual results were combined via higher-level analysis using fixed-effects modeling ^60^. In this step, the individual results were registered to the SIGMA rat brain atlas as a common reference space^61^. The resulting group-level Z statistic images were then thresholded using voxelwise inference with the Gaussian random field (GRF) method for family-wise error (FWE) correction with a threshold of p = 0.01^62^.

fMRI activation volume of separate regions was quantified by calculating the proportion of activated voxels relative to the total number of voxels in the respective regions of the SIGMA brain atlas, expressed as ‘fraction of region activated’ and plotted as mean ± one standard error of the mean (Figure S2A-C). To calculate the total activated volume, the number of all activated voxels was multiplied by the voxel volume. Volume differences along different frequency-stimulation parameters were assessed using one-way ANOVA p <0.05 with Tukey-Kramer multiple comparisons correction.

Differences in fMRI activation between groups (e.g. PL vs IL) were analyzed using two-sample unpaired t-tests (see the user guide of the FSL function “FEAT”^63^) and an FWE-corrected voxelwise significance threshold of p = 0.01.

## Acknowledgments

This work was supported by the BrainLinks-BrainTools Center, funded by the Federal Ministry of Economics, Science and Arts of Baden-Württemberg within the Excellence Initiative II; German Research Foundation (DFG, LE 3437/2-1, DI1908/11-1, DI1908/7-1, DI1908/6-1, DI1908/12-1, DI1908/14-1); BMBF Bernstein Prize for Computational Neuroscience 01GQ1301, Core Facility Advanced Molecular Imaging Research Center, Department of Radiology – Medical Physics of the University Hospital Freiburg; and EU-project euSNN (MSCA-ITN-ETN H2020-860563). We extend our sincere gratitude to Jan Deubner and Dominic Lau for assisting in training, surgery, and optogenetic measurements. We are very thankful to Lidia Miguel Telega and Máté Döbrössy for the use of their facilities. We would also like to express our appreciation to Enya Paschen and Carola Haas for their assistance in sharing their surgical techniques for fMRI in rodents, allowing us to assess various approaches for such experiments.

## CRediT authorship contribution statement

**Z.J.:** Conceptualization, Methodology, Validation, Formal analysis, Investigation, Data Curation, Writing - Original Draft, Writing - Review & Editing, Visualization. **N.S.:** Conceptualization, Methodology, Software, Validation, Formal analysis, Investigation, Data Curation, Writing - Review & Editing, Visualization. **A.A.:** Conceptualization, Methodology, Formal analysis. **F.S.:** Conceptualization, Formal analysis, Visualization. **S.H.:** Methodology, Investigation. **K.F.:** Formal analysis, visualization. **C.B.:** Conceptualization, Formal analysis, visualization. **M.Z.:** Conceptualization, Resources, Writing - Review & Editing, Funding acquisition. **I.D.:** Conceptualization, Methodology, Resources, Writing - Review & Editing, Supervision, Project administration, Funding acquisition.

## Supplemental figures

**Supplementary Figure 1:**
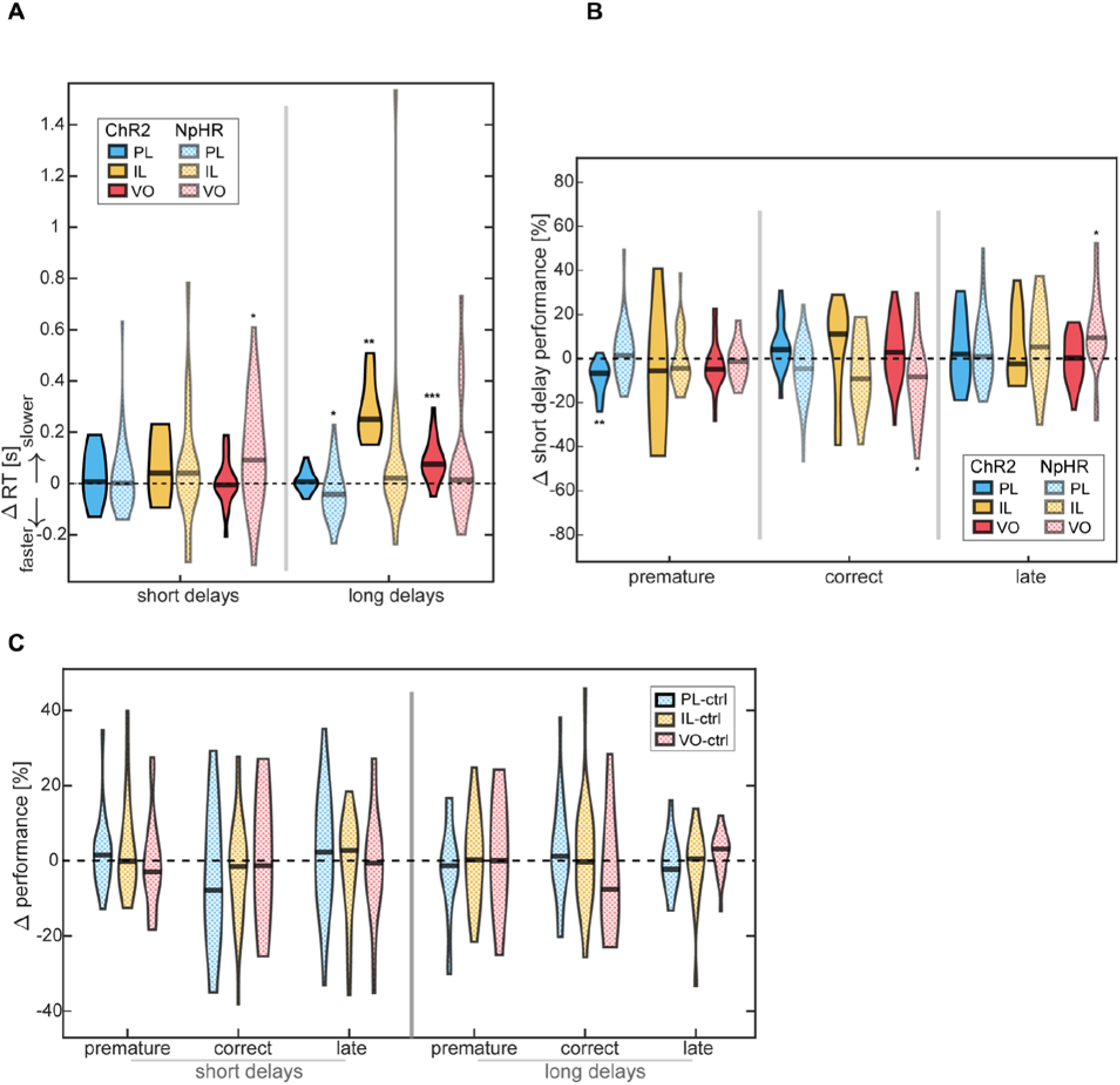
Optogenetic behavioral effects. 30 Hz blue light stimulation effects in the PL (blue), IL (yellow), and VO (pink); effects from previously published *Hardung et al*. data for each corresponding subarea inhibited via continuous yellow light illumination of NpHR-injected rats plotted in lighter shades. Median value: black bar. (*) p <0.0083, (**) p <0.0017, (***) p <0.00017 (Wilcoxon signed-rank test, Bonferroni-corrected p-thresholds **(A)** Difference in reaction times (RT) (laser vs. no-laser). Correct and late trial RTs (time to release after the tone presentation) are pooled in each session to acquire a difference in mean RT per session. Shape widths indicate the distribution of differences between mean RT laser vs. no-laser (in each outcome type) per session. See also Table S3. **(B)** Difference (laser vs. no-laser) in session-wise performance of short-delay trials over all sessions in each cohort. Shape widths indicate session-wise effect distribution. See also Table S1. **(C)** Behavioral blue-light control experiments conducted in NpHR-injected and implanted rats. Difference (laser vs. no-laser) in session-wise performance; median indicated by black bar; shape widths indicate session-wise effect distribution. Continuous blue light (10–12 mW) did not produce any significant effects between laser and non-laser stimulated trials. Rats (N) and sessions (n) for each group: PL (N=4, n=22), IL (N=3, n= 28), VO (N=3, n=19).

**Supplementary Figure 2:**
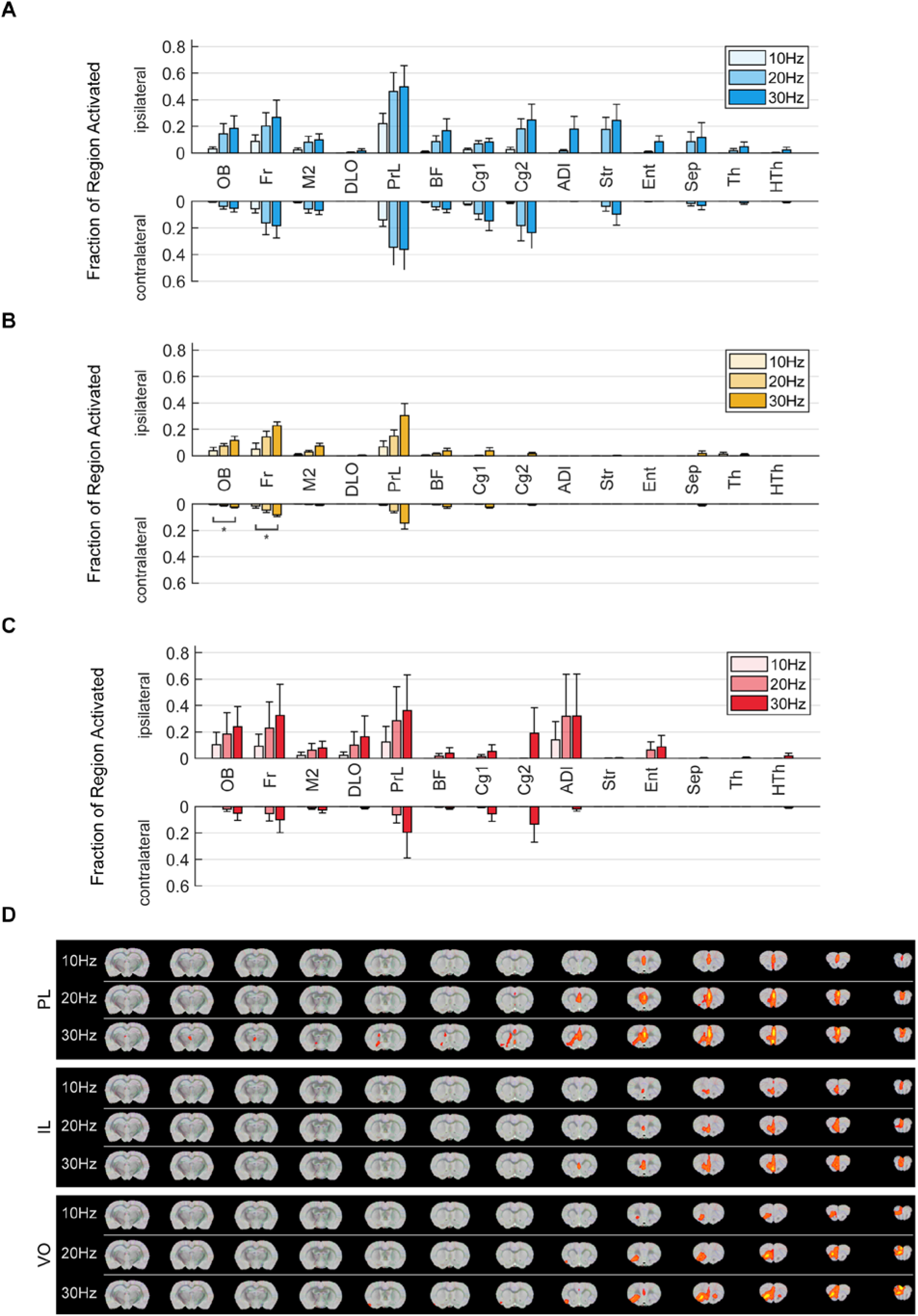
Activation area from stimulation of individual PFC regions. Proportion (range [0–1]) of the activated volume of individual recruitment areas in the ipsi- and contralateral hemisphere to the implanted fiber and light stimulation. **(A)** PL stimulation revealed fMRI activation near the fiber tip, extending from the medial part of the prelimbic area (PrL) to the primary cingulate cortex (Cg1) and olfactory bulb (OB), with 10 Hz stimulation yielding limited activation volume. Higher stimulation frequencies (20–30 Hz) expanded activation areas, with similar ipsi- and contralateral recruitment of the frontal associated cortex (Fr), the secondary motor cortex (M2), PrL, and primary and secondary cingulate cortex (Cg1, Cg2). There was additional ipsilateral recruitment at 20–30 Hz of the OB, basal forebrain region (BF), striatum (Str), and septum (Sep) and at 30Hz of the agranular dysgranular insular cortex (ADI), entorhinal cortex (Ent) and thalamus (Th). Overall, a trend (n.s.) indicated a larger total activation volume with increasing frequency. **(B)** IL stimulation evoked limited fMRI activation, mainly constricted to the fiber tip area, gradually expanding with higher-frequency stimulation. At 10–20 Hz stimulation, activity was observed in the PrL, OB, and medial Fr. 30 Hz activated the same areas, with additional recruitment of M2. There was a significantly larger total contralateral area activated with 30 Hz, specifically in the contralateral recruitment of OB and Fr (compared to 10 Hz). **(C)** VO stimulation activated the fiber tip region in the ventrolateral part of the PrL and part of the OB, along with M2 and Fr. Lower frequencies revealed low activation volume in PrL, OB, Fr, and ADI exclusively ipsilateral to the fiber; a contralateral response could only be measured from 20 Hz onwards, activating the same areas as 10 Hz. Contralateral recruitment involved the Fr and PrL, and the additional recruitment of Ent and the dorsolateral orbital cortex (DLO). 30 Hz activated all lower-frequency areas alongside Cg1 and Cg2. Overall, a trend (n.s.) indicated a larger total activation volume with increasing frequency. **OB**: Olfactory Bulb, **Fr**: Frontal Associated Cortex, **M2**: Secondary Motor Cortex, **DLO**: Dorso Lateral Orbital Cortex, **PrL**: Prelimbic System, **BF**: Basal Forebrain Region, **Cg1**: Primary Cingulate Cortex, **Cg2**: Secondary Cingulate Cortex, **ADI**: Agranular Dysgranular Insular Cortex, **Str**: Striatum, **Ent**: Entorhinal Cortex, **Sep**: Septal Region, **Th**: Thalamus, **HTh**: Hypothalamus. (*) p *<*0.05, (**) p *<*0.01, (***) p *<*0.001 (one-way ANOVA, Tukey-Kramer multiple comparison). **(D)** Group-level activation maps for PL, IL, and VO cohorts across 10, 20, and 30 Hz stimulation.

**Supplementary Figure 3:**
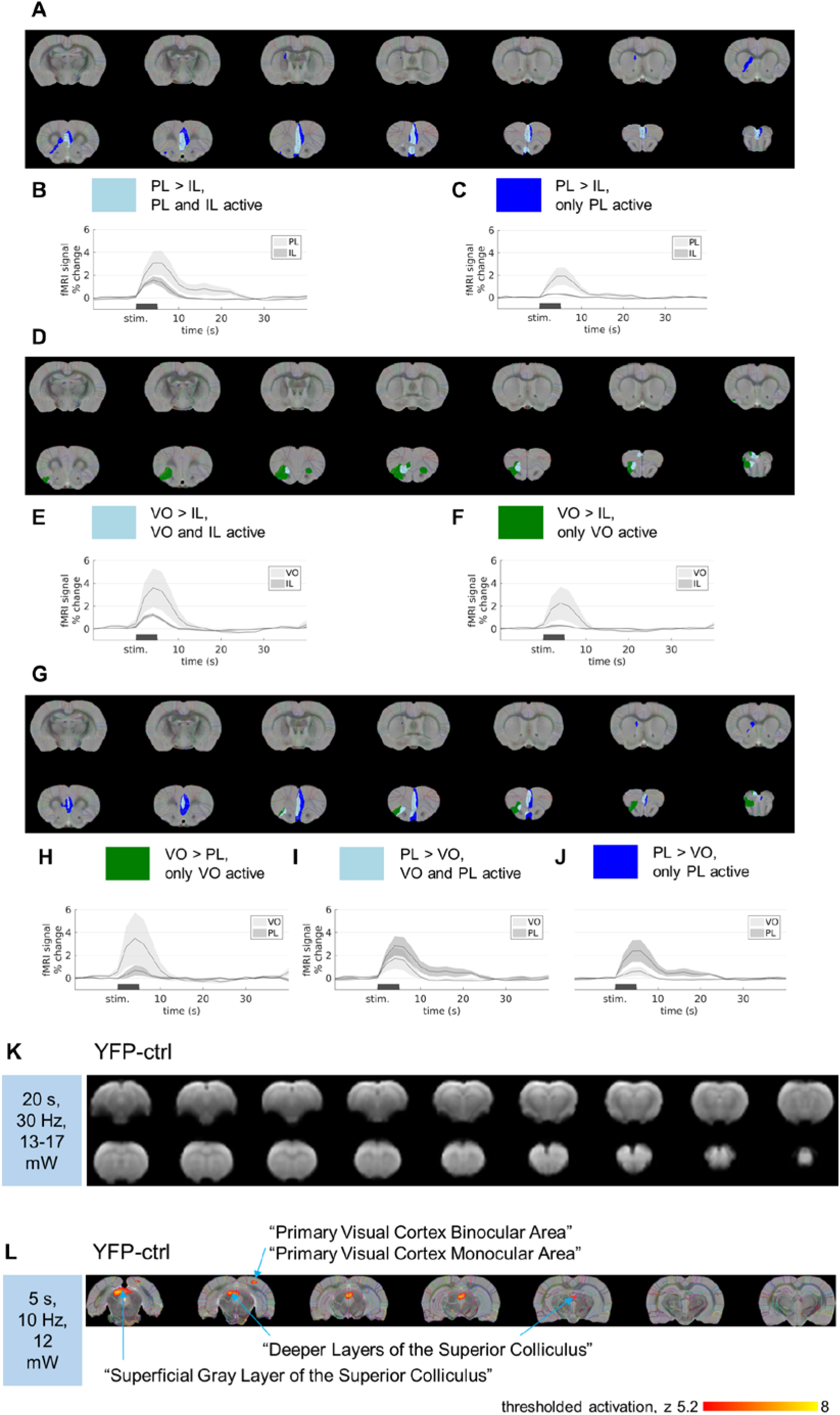
Activation-intensity differences between PFC subsections. **(A-J)** Maps illustrate activation-intensity differences between two PFC groups during 30 Hz stimulation (two-sample unpaired t-tests, FWE-corrected voxelwise significance threshold of p = 0.01); activated voxels are highlighted in color. Mean fMRI response is depicted in time series plots with shaded regions representing SEM. Differences between two groups (based on the activation maps, Figure S2D) derive from one group displaying activation and the other not, or from both exhibiting activation while one group has greater activation strength. Three PL, four IL, and three VO rats are compared group-wise. **(A-C)** PL vs. IL stimulation. **(A)** In the colored areas (light blue and dark blue), PL stimulation created a significantly stronger response than IL stimulation. No areas were more strongly activated by IL stimulation than PL stimulation. **(B)** The mean time series for light blue areas (the fiber tip-surrounding area in the prelimbic area (PrL) and extending into the olfactory bulb (OB), primary cingulate cortex (Cg1), basal forebrain (BF) and frontal associated cortex (Fr)), indicating commonly activated areas during PL and IL stimulation, where PL stimulation resulted in stronger activation. **(C)** The dark blue area represents the activated area, including the striatum (Str), which responds exclusively to PL stimulation, illustrated in the mean time series. **(D-F)** VO vs. IL stimulation. In the colored areas (light blue and green), VO stimulation had a significantly stronger response than IL stimulation. No areas were more strongly activated by IL stimulation than by VO stimulation. **(E)** Light blue shaded areas depict co-activated areas (the ventrolateral PrL and anteromedial parts of Fr and OB), where VO stimulation was stronger. **(F)** The time series for the green area illustrates more lateral areas, reaching the entorhinal cortex (Ent), that were activated by VO stimulation, but not IL stimulation. **(G-J)** PL vs. VO stimulation. Colored areas represent significant differences in the responses induced by PL and VO stimulation. **(H)** The mean time series for green areas that were activated exclusively by VO stimulation, including an area extending from the PrL in the ventrolateral direction and anterolateral areas of the OB, Fr, and DLO. **(I)** The mean time series for light blue areas reveals co-activation by PL and VO, but slightly stronger activation from PL stimulation in a region along the dorso-ventral axis; this difference is relatively weak since both PL and VO stimulation showed an fMRI response. **(J)** The mean time series for dark blue areas shows activation only from PL stimulation, extending along the dorso-ventral axis, additionally including parts of the Str. **(J-K)** Blue light control opto-fMRI observations. **(K)** Blue light stimulation in PL targeted YFP control animals. Group-level fMRI map from blue light stimulation for 20 s blocks (2 ms-wide pulses, 30 Hz, 13–17 mW), averaged from three datasets between two animals, also representative of results for 5 s block stimulation. Slices are displayed in a caudal to rostral direction (top left to bottom right) of the mean functional image, registered to the reference space. Any fMRI activation would be overlaid in color, however no response was detected at a FWE-corrected voxelwise significance threshold of p = 0.01. This indicates that there was no significant heating, which would be observable by fMRI responses around the fiber tip. **(L)** A single subject result showing visual activation in the caudal part of the brain, registered into reference space. fMRI activation from blue light stimulation (5 s blocks at 10 Hz and 12 mW) in a YFP-injected, PL-implanted rat is shown in red-yellow (the color represents the z-score from 5 (red) to 8 (yellow) at a FWE-corrected voxelwise significance threshold of p = 0.01). While visual activation was not consistently observed, this dataset was selected to illustrate a case of visual activation. Lack of response around the fiber tip indicates that heating effects are negligible. Occasionally observed activation was confined to areas responsible for visual processing^64–66^, specifically the superior colliculus and visual cortex^64–66^. These areas were not relevant in our study and therefore were not considered.

**Supplementary Figure 4:**
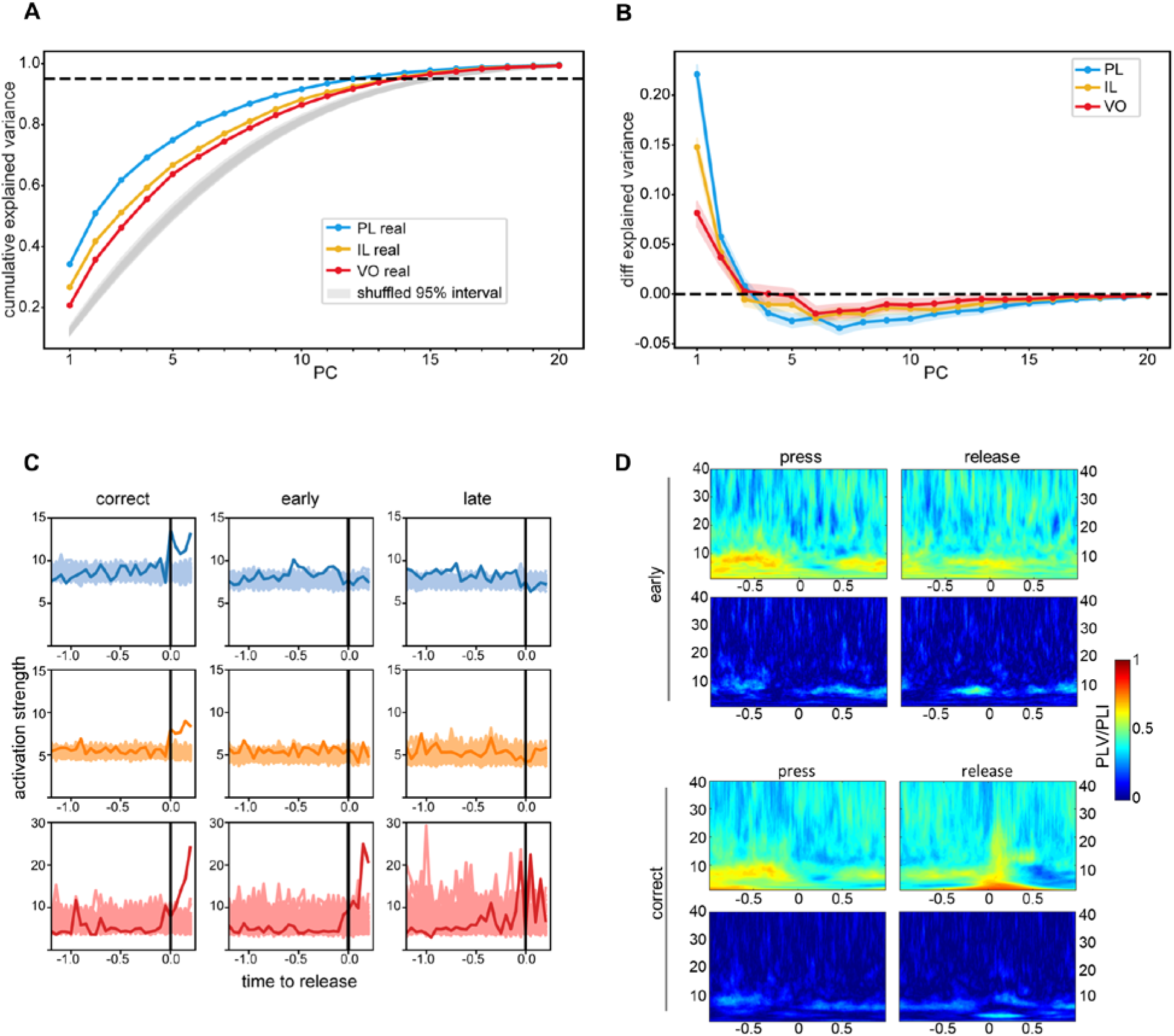
Supplementary results of PFC neural recording during the response-preparation task. **(A)** Cumulative explained variance ratios per PC for each PFC subsection compared to random cumulative explained variance per PC (there was no significant difference between the shuffled datasets; the 95^th^ percentiles of these datasets shown as grey shaded area). Dashed line indicates 95% threshold of explained variance. **(B)** Difference in explained variance between each real PC and the corresponding PC in each shuffled dataset, with the mean as a dark line and 95^th^ percentile as the lightly shaded region. Dashed line indicates cutoff for significant PC selection, where explained variance per PC from the shuffled data is greater than that from the real data. **(C)** Average activation strength (bold line) aligned to press (t=0) over all correct (left column), early (middle column), and late trials (right column) for simultaneously recorded activity from PL (blue), IL (yellow), or VO (pink) neurons. Activation strength from time-shuffled data is plotted in a lighter shade. **(D)** PLV (top row) and PLI (bottom row) spectrograms from single-animal trial-averaged PL-IL LFP signals in correct (right) and premature (left) trials, time-aligned to press (1^st^ and 3^rd^ column) or release (2^nd^ and 4^th^ column). Y-axis: frequency (Hz), x-axis time in seconds. Color bar indicates the PLV or PLI for all maps using the same scale.

## Supplemental tables

**Supplementary Table 1:** 30 Hz stimulation effect on behavioral performance. ChR2 stimulation results for this experiment, and reanalyzed data from Hardung et. al. from NpHR inhibition experiments, corresponding to Figure 1D, as well as corresponding values in the short-delay trials (Figure S1B). Median values for the difference in error rate between laser and non-laser trials, along with the 1^st^ and 3^rd^ Quartile, and IQR (interquartile range). P-values from Wilcoxon signed-rank test between and session-wise error rates of laser- and non-laser-stimulated trials, for each cohort. (*) p <0.0083, (**) p <0.0017, (***) p <0.00017 (Wilcoxon signed-rank test, Bonferroni-corrected p-thresholds). Rats (N) and sessions (n) for each ChR2 cohort: PL (N=3, n=17), IL (N=3, n=12), and VO (N=4, n= 28); rats and sessions from Hardung et al. experiments: PL (N=4, n=44), IL (N=3, n=27), VO (N=4, n =23).

|  | Trial type | Trial-Outcome | Area | Median | 1<sup>st</sup>Quartile | 3<sup>rd</sup> Quartile | IQR | p-value |
| --- | --- | --- | --- | --- | --- | --- | --- | --- |
| ChR2-pulsed blue light 30 Hz | short delay | premature | PL | -6.67 | -13.04 | -4.39 | 8.65 | 0.0005** |
|  |  |  | IL | -5.54 | -41.11 | 3.91 | 45.02 | 0.2036 |
|  |  |  | VO | -4.90 | -8.29 | 0.93 | 9.23 | 0.0679 |
|  |  | correct | PL | 4.00 | -0.83 | 10.70 | 11.52 | 0.0554 |
|  |  |  | IL | 11.13 | -8.82 | 20.63 | 29.44 | 0.4697 |
|  |  |  | VO | 2.75 | -6.64 | 11.40 | 18.04 | 0.2645 |
|  |  | delayed | PL | 2.00 | -10.18 | 15.67 | 25.85 | 0.5540 |
|  |  |  | IL | -2.34 | -7.68 | 20.00 | 27.68 | 0.5693 |
|  |  |  | VO | 0.25 | -6.62 | 9.29 | 15.90 | 0.7156 |
|  | long delay | premature | PL | -22.30 | -41.44 | -12.28 | 29.16 | 0.0008** |
|  |  |  | IL | -79.99 | -89.07 | -12.43 | 76.64 | 0.0024* |
|  |  |  | VO | -27.57 | -36.72 | -18.40 | 18.32 | 0.00001*** |
|  |  | correct | PL | 24.29 | 12.98 | 41.47 | 28.49 | 0.0007** |
|  |  |  | IL | 60.38 | 12.43 | 66.97 | 54.54 | 0.0020* |
|  |  |  | VO | 21.63 | 13.69 | 31.45 | 17.77 | 0.00004*** |
|  |  | delayed | PL | -1.16 | -2.25 | 0.00 | 2.25 | 0.2783 |
|  |  |  | IL | 14.31 | 0.00 | 27.34 | 27.34 | 0.0078* |
|  |  |  | VO | 5.01 | 0.00 | 8.71 | 8.71 | 0.0007** |
| NpHR- yellow continuous light | short delay | premature short | PL | 1.48 | -6.37 | 8.84 | 15.21 | 0.3683 |
|  |  |  | IL | -4.48 | -9.11 | 11.18 | 20.29 | 0.5338 |
|  |  |  | VO | -1.35 | -7.14 | 4.36 | 11.50 | 0.4077 |
|  |  | correct short | PL | -4.66 | -13.35 | 4.47 | 17.82 | 0.0184 |
|  |  |  | IL | -9.13 | -15.81 | 7.70 | 23.51 | 0.1068 |
|  |  |  | VO | -8.31 | -21.63 | -2.86 | 18.77 | 0.0081* |
|  |  | delayed short | PL | 0.88 | -7.97 | 12.08 | 20.04 | 0.3327 |
|  |  |  | IL | 5.30 | -5.33 | 21.15 | 26.48 | 0.1299 |
|  |  |  | VO | 9.40 | 2.42 | 19.70 | 17.28 | 0.0024* |
|  | long delay | premature long | PL | 7.10 | -0.33 | 20.23 | 20.56 | 0.0004** |
|  |  |  | IL | -13.10 | -16.81 | -0.57 | 16.24 | 0.0005** |
|  |  |  | VO | -3.04 | -14.41 | 10.49 | 24.89 | 0.5430 |
|  |  | correct long | PL | -8.52 | -17.10 | -0.53 | 16.57 | 0.0003** |
|  |  |  | IL | 7.14 | -0.90 | 13.47 | 14.37 | 0.0116 |
|  |  |  | VO | -2.62 | -19.34 | 7.66 | 27.00 | 0.5430 |
|  |  | delayed long | PL | -1.55 | -4.02 | 3.43 | 7.45 | 0.2990 |
|  |  |  | IL | 4.10 | -1.41 | 8.82 | 10.22 | 0.0174 |
|  |  |  | VO | 2.73 | -0.54 | 6.40 | 6.94 | 0.1080 |

**Supplementary Table 2:** Blue light stimulation frequency-dependent effects on long premature error rate. Values and statistics for % change of premature error rate in long trials for different stimulation-frequency and control parameters (yellow light which is non-activating for ChR2), including data corresponding to Figure 1E. Median values for the difference in error rate between laser and non-laser trials, along with the 1^st^ and 3^rd^ Quartile, and IQR (interquartile range). A positive value indicates that the error rate increased during stimulation in comparison to no stimulation. One-way ANOVA was performed for each cohort to check for differences in the laser effect on trial-outcome error rate among the stimulation frequencies and control parameters. If the null hypothesis was rejected, multicomparison with Tukey-Kramer was performed between the stimulation parameters to acquire the given p-values. (*) p <0.05, (**) p <0.01, (***) p <0.001-groups for which the ANOVA did not reject the null hypothesis identified as n.s. (non-significant). P-values given are for comparison to the same cohort under control parameters. The last column shows any other significant p-values between various frequency-stimulation parameters within the same cohort (these are illustrated with blue stars for the PL cohort and red stars for the VO cohort in Figure 1E). Data from 3 PL, 3 IL, and 4 VO rats over several sessions (n) per stimulation parameter. PL/Ctrl: n=22; PL/10 Hz: n=25; PL/20 Hz: n=58; PL/30 Hz: n=17. IL/Ctrl: n=13; IL/10 Hz: n=17; IL/20 Hz: n=12; IL/30 Hz: n=12. VO/ctrl: n=23; VO/10 Hz: n=20; VO/20 Hz: n=29; VO/30 Hz: n=28.

| Area | Stimulation parameters | Median | 1<sup>st</sup>Quartile | 3<sup>rd</sup> Quartile | IQR | p-value (difference to control) | p-value (difference between stimulation frequencies) |
| --- | --- | --- | --- | --- | --- | --- | --- |
| PL | ctrl | 0.38 | -4.12 | 11.32 | 15.44 | -- | -- |
|  | 10 | -14.58 | -19.42 | -5.99 | 13.43 | 0.0016** | -- |
|  | 20 | -11.02 | -26.58 | -3.53 | 23.05 | 0.0002*** | 30 Hz: 0.029857* |
|  | 30 | -22.30 | -41.44 | -12.28 | 29.16 | 4.7928e-07*** | 20 Hz: 0.029857* |
| IL | ctrl | 0.75 | -1.32 | 4.08 | 5.40 | -- | -- |
|  | 10 | -27.78 | -42.75 | -11.72 | 31.03 | 0.06921 | 30 Hz: 0.0225* |
|  | 20 | -58.61 | -82.30 | -3.85 | 78.45 | 0.00072*** | -- |
|  | 30 | -79.99 | -89.07 | -12.43 | 76.64 | 3.4245e-05*** | 10 Hz: 0.0225* |
| VO | ctrl | 1.13 | -9.44 | 6.51 | 15.95 | -- | -- |
|  | 10 | -3.48 | -8.99 | 6.60 | 15.58 | 0.8385 | 20 Hz: 0.0145*, 30 Hz: 0.0007*** |
|  | 20 | -25.42 | -34.42 | -9.47 | 24.96 | 0.00048*** | 10 Hz: 0.0145* |
|  | 30 | -27.57 | -36.72 | -18.40 | 18.32 | 1.144e-05*** | 10 Hz: 0.0007*** |

**Supplementary Table 3:**
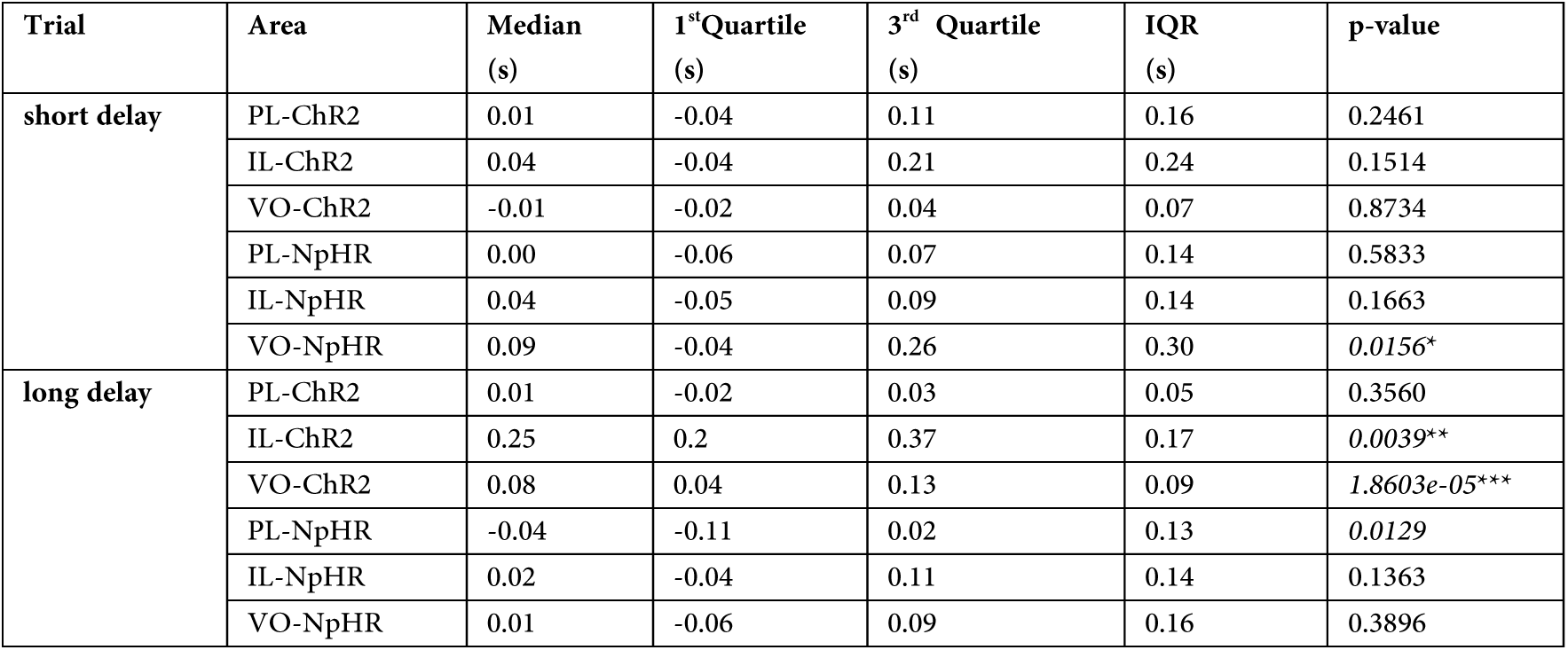
30 Hz stimulation effects on reaction time. Data corresponding to Figure S1A. Median, 1st and 3^rd^ Quartile, and IQR (interquartile range) values for the session-wise difference in mean reaction time between laser and non-laser trials for short or long-delay trials in which the lever release occurred after the tone (correct and late trials pooled for each session). P-values from Wilcoxon signed-rank test between mean reaction times of laser-stimulated and non-laser-stimulated trials per session, for each cohort. Minus signs refer to a reduction of RTs upon laser stimulation. Unadjusted p-values are reported in the last column: (*) p <0.0250, (**) p <0.0050, (***) p <3.33e-04 (Wilcoxon signed-rank test, Bonferroni-corrected p-thresholds). Rats (N) and sessions (n) for each ChR2 cohort: PL (N=3, n=17), IL (N=3, n=12), and VO (N=4, n=28).

